# Multi-omics characterization of interaction-mediated control of human protein abundance levels

**DOI:** 10.1101/499434

**Authors:** Abel Sousa, Emanuel Gonçalves, Bogdan Mirauta, David Ochoa, Oliver Stegle, Pedro Beltrao

## Abstract

Proteogenomic studies of cancer samples have shown that copy number variation can be attenuated at the protein level, for a large fraction of the proteome, likely due to the degradation of unassembled protein complex subunits. Such interaction mediated control of protein abundance remains poorly characterized. To study this we compiled genomic, (phospho)proteomic and structural data for hundreds of cancer samples and find that up to 42% of 8,124 analyzed proteins show signs of post-transcriptional control. We find evidence of interaction dependent control of protein abundance, correlated with interface size, for 516 protein pairs, with some interactions further controlled by phosphorylation. Finally, these findings in cancer were reflected in variation in protein levels in normal tissues. Importantly, expression differences due to natural genetic variation were increasingly buffered from phenotype differences for highly attenuated proteins. Altogether, this study further highlights the importance of post-transcriptional control of protein abundance in cancer and healthy cells.

## Introduction

Cancer cells can harbor large number of somatic DNA alterations ranging from point mutations to gene copy changes that can occur from deletion or amplification of small regions or whole chromosomes. While these events are the source of the genetic variation that can confer a selective advantage and lead to cancer, large changes in gene numbers can be detrimental and cause imbalances in the corresponding protein levels. Several studies have shown that the majority of changes in gene copy number will propagate to changes in the corresponding protein levels (Dephoure et al., 2014; Pavelka et al., 2010; Stingele et al., 2012). However, models of aneuploidy of different species and analysis of gene copy number variation (CNV) in cancer have shown that CNVs of protein coding genes belonging to protein complexes tend to be attenuated at protein level (Dephoure et al., 2014; Gonçalves et al., 2017; Ishikawa, Makanae, Iwasaki, Ingolia, & Moriya, 2017). In addition we have shown that some complex members can act as rate-limiting subunits and indirectly control the degradation level of attenuated complex members (Gonçalves et al., 2017). These results are in-line with pulse chase degradation measurements showing that several complex subunits have a two-state degradation profile, that is compatible with a model in which they are expressed above the required levels and have a higher degradation rate when unbound from the complex (McShane et al., 2016). The attenuation of changes at the protein level also justifies why protein complex subunits show higher correlation of protein abundances than the corresponding mRNA levels (Ryan, Kennedy, Bajrami, Matallanas, & Lord, 2017; Wang et al., 2017) and why correlation analysis can be used to identify cancer specific interaction networks (Lapek et al., 2017; Roumeliotis et al., 2017).

These results support a long standing view that protein complex formation can set the total amount of protein levels (Abovich, Gritz, Tung, & Rosbash, 1985). The degradation of unbound subunits may be due to a requirement of avoiding free hydrophobic interface surfaces that can be prone to aggregate (Young, Jernigan, & Covell, 1994). In eukaryotic species this appears to be achieved by degrading excess production while in bacterial and archaeal species genes coding for protein complexes subunits tend to occur within operon structures such that they will be expressed at similar levels (Mushegian & Koonin, 1996). This link between appropriate expression and complex formation is further emphasized by the preferential ordering of subunits in operons starting from the subunits that tend to assemble first (Wells, Bergendahl, & Marsh, 2016).

While this phenomenon of gene dosage attenuation in protein complexes has been well documented we still do not understand (i) what protein properties are associated with the propensity for a protein to be attenuated, (ii) nor if the characteristics of the attenuation process are seen in non cancerous cells. Here we have extended on a previous analysis (Gonçalves et al., 2017), performing a multi-omics study of protein level attenuation of gene dosage that combines genomics, (phospho)proteomics and structural data. Analysing 8,124 genes/proteins we observed that up to 42% of proteins show evidence of post-transcriptional regulation. Over 500 protein-protein interactions show indirect control of degradation of one subunit via physical associations, 32 of which may be further controlled by phosphorylation. Using structural models for 3,082 interfaces we find that a higher fraction of interface residues is associated with a higher degree of attenuation. Finally, we studied the impact of these findings on non-cancerous systems. We find that protein interaction mediated control of protein abundances have an impact of the variation of protein levels across tissues and that the degree of attenuation correlates with the probability that natural variation with an impact on gene expression may result in a phenotypic consequence.

## Results

### Protein level attenuation of gene dosage associates with distinct essentiality and structural features

In order to study protein post-transcriptional control we collected matched gene copy-number, mRNA and protein expression cancer datasets made available by TCGA and CPTAC consortia, for breast (BRCA) (Cancer Genome Atlas Network, 2012b; Mertins et al., 2016), ovarian (HGSC) (Cancer Genome Atlas Research Network, 2011; H. Zhang et al., 2016) and colorectal (COREAD) cancers (Cancer Genome Atlas Network, 2012a; B. Zhang et al., 2014). In addition we compiled existing protein/gene expression and copy-number data for cancer cell lines from *Lapek et al*. (BRCA) (Lapek et al., 2017), *Roumeliotis et al*. (COREAD) (Roumeliotis et al., 2017) and *Lawrence et al*. (BRCA) (Lawrence et al., 2015). In total, 368 cancer samples (294 tumours and 74 cell lines) were compiled in our study with matched gene expression, copy-number and protein abundance (**Figure 1A**). Principal component analysis (PCA) revealed the presence of confounding effects in the RNA and protein expression data (**Figures S1A and S2A**). These effects are related to cancer type, experimental batch, type of proteomics experiment, and also patient gender and age. Therefore, these potential confounding effects were regressed-out from the RNA and protein expression data (**Methods**). After correction, the association between the principal components and the potential confounding effects was removed (**Figures S1B and S2B**).

**Figure 1.**
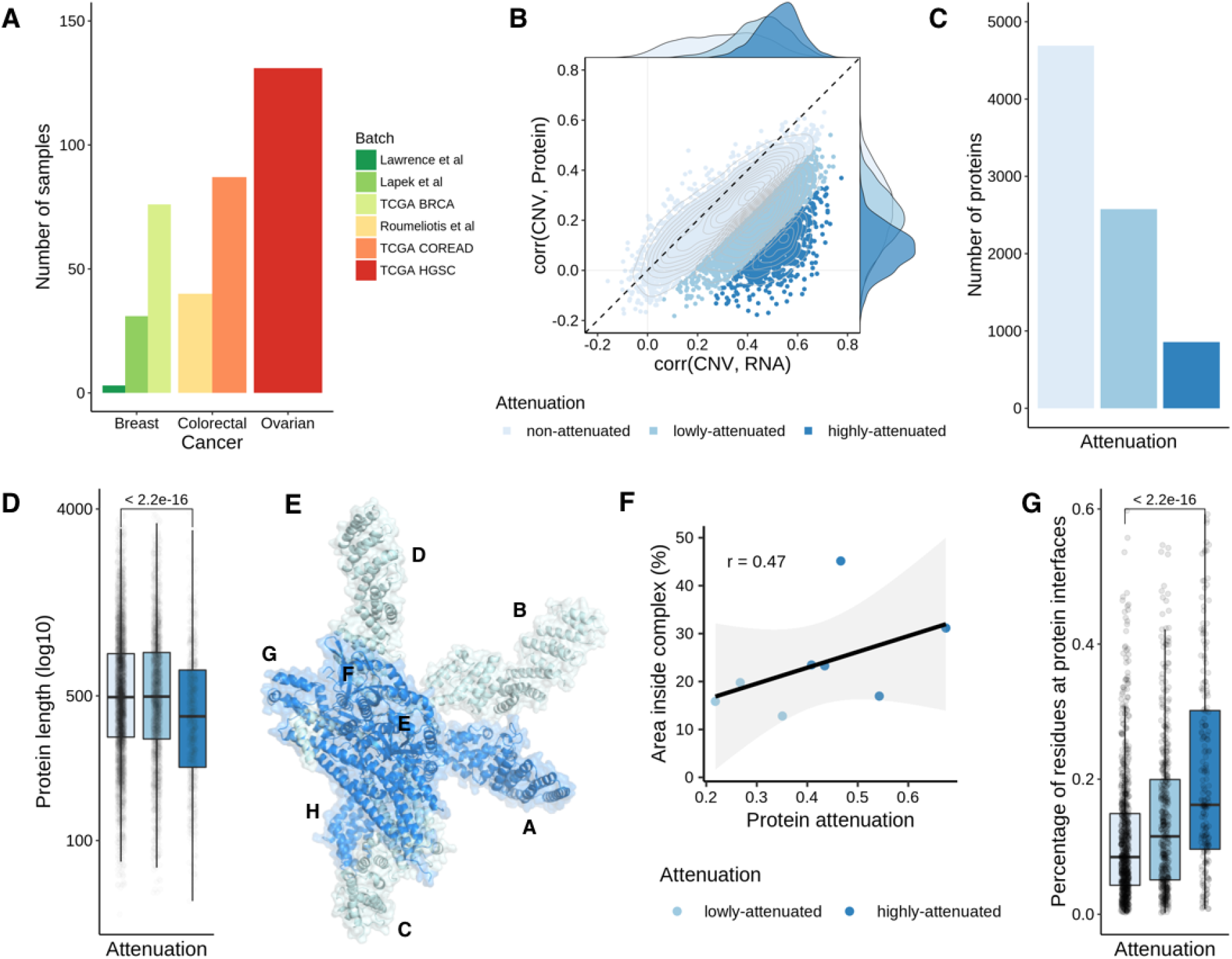
Features of proteins showing gene dosage buffering at the protein level. **(A)** Number of samples with CNV, mRNA and protein measurements, by cancer type and batch. **(B)** Scatter plot representing the correlation between the CNV and mRNA (x-axis) and the CNV and protein (y-axis), for each gene. The colors represent the attenuation levels. From light blue to dark blue: non-attenuated, lowly-attenuated and highly-attenuated. **(C)** Number of proteins by attenuation level. **(D)** Protein length (log10 of number of residues) by attenuation level. **(E)** Representation of COP9 signalosome complex. **(F)** Scatter plot representing the correlation between the attenuation potential (x-axis) and the fraction of residues at interfaces in the complex (y-axis), for the 8 protein subunits from the COP9 signalosome complex represented in figure E. **(G)** Percentage of residues at protein interfaces by attenuation level.

We then investigated the impact of CNV in cancer proteomes, using the strategy reported in *Gonçalves et al*. (Gonçalves et al., 2017). Due to the sparseness of the protein data we selected genes with protein measurements in at least 25% of the 368 samples, comprising 8,124 genes with CNV, mRNA and protein expression. For each gene, we then calculated the Pearson correlation coefficient between the CNV and the mRNA and the CNV and the protein, across samples. In order to assess the disagreement between the transcriptome and proteome regarding the copy number changes, we calculated an attenuation potential, corresponding to the difference between Pearson coefficients (**Methods**). A higher attenuation potential suggests genes that have CNVs buffered at the protein level. As previously, we then clustered the genes by attenuation potential using an unsupervised gaussian mixture model. Using this strategy, we identified 3,435 (42%) genes as attenuated at the protein level (2,578 low-attenuated and 857 high-attenuated) and 4,689 as non-attenuated (**Figure 1B** and **1C** and **Supplementary Table 1**). These results indicate that up to 42% of genes show signs of gene dosage buffering at the protein level, probably due to a post-transcriptional control of protein degradation, and robustly recapitulates previous findings on a smaller set of 6,418 genes (Gonçalves et al., 2017). In line with previous findings, the list of attenuated genes is strongly enriched in well characterized protein complex members, and notably in members of large complexes (**Figure S3**). More, the attenuation potential is correlated with the number of subunits in a protein complex, indicating that members of large complexes have higher attenuation than those of small complexes (**Figure S3E**). Attenuated genes are also expected to show increased ubiquitination after proteasome inhibition, which was confirmed here using previously published data with 3 different proteasome inhibitors – MG-132, epoxomicin and bortezomib (Higgins et al., 2015; W. Kim et al., 2011; Udeshi et al., 2013; S. A. Wagner et al., 2011) (**Figure S4A**). Having defined a comprehensive list of genes/proteins with different degrees of attenuation we then set out to characterize their physical and genetic properties.

We first asked if the level of attenuation relates to distinct essentiality features, based on gene essentiality defined by CRISPR-Cas9 screens (Meyers et al., 2017). Highly-attenuated proteins showed higher gene essentiality than low and non-attenuated proteins (**Figure S4B**) (Wilcoxon rank-sum test p-value < 2.2e-16, highly– vs non-attenuated proteins). This result is likely to be driven by the enrichment of protein complex members of essential complexes, such as the ribosome and spliceosome. We then studied the physical characteristics of these proteins such as length and structural properties. We found that the highly-attenuated proteins tend to have a smaller size (**Figure 1D**) (Wilcoxon rank-sum test p-value < 2.2e-16; highly– vs non-attenuated proteins), suggesting a size-dependent buffering mechanism. For the structural analysis, we considered a total of 2,392 proteins having structurally defined interface models (Mosca, Céol, & Aloy, 2012). We illustrate this analysis with the COP9 signalosome complex (**Figure 1E**) where we noticed a trend in which the subunits with a larger surface buried in interfaces had the strongest attenuation (**Figure 1F**). This trend was seen across all proteins, with the average fraction of residues at interfaces increasing from the non-attenuated to the highly-attenuated proteins (**Figure 1G**).

### Protein interaction-dependent control of degradation depends on interface size

The features of highly attenuated proteins suggest that protein interactions are an important determinant of a protein’s susceptibility of having gene dosage attenuation. It has been suggested that some members of protein complexes can act as scaffolding or rate-limiting subunits. We have previously analyzed a set of 58,627 protein interactions among complexes curated in CORUM database and identified a set of 48 interactions in which a protein can indirectly control the abundance of an interacting partner (Gonçalves et al., 2017). Here we set out to expand this analysis to all currently reported human physical interactions in the BioGRID database (**Methods**). In total, we collected 572,856 physical interactions and identified proteins whose CNV changes correlate with the protein abundance of interacting proteins once their mRNA levels are taking into account (**Methods**). For an interaction pair of proteins X and Y, we used a linear regression model, where we predict the protein levels of protein Y using the CNV of X, discounting the mRNA of Y and the impact of other covariates (**Methods**). Correlating molecular changes with DNA variation such as CNVs ensures the correlations found are most likely causal and in the direction of DNA changes to the molecular changes. Copy number alterations in cancer most often occur in large segments leading to co-amplification or co-deletion of multiple co-localized genes. For proteins with two or more interacting partners that are genomically co-localized, we selected only the top ranking association to avoid spurious “passenger” associations (**Methods**).

Out of 572,856 physical interactions we had data to test associations for 411,591 with this model, finding 516 protein-protein associations as significant using CNV and mRNA (FDR < 5%) (**Figure 2A** and **Supplementary Table 2**). In this set of associations, we classified the proteins as *controlling* (423) – those capable of controlling the protein levels of their interactions partners; *controlled* (353) – whose abundance levels depends on their interactions; and *both* (60), as the proteins with the two characteristics (**Figure 2B**). Out of 423 *controlling* proteins, 62 had at least two interactions. The top *controlling* protein was TCP1, which was predicted to control the protein abundance of 7 complex partners, including CCT3, CCT5, CCT7 and CCT8 (**Figure 2D**). As expected, the *controlled* proteins had higher attenuation potential, a consequence of the post-transcriptional regulation of their protein levels (**Figure 2C**) (Wilcoxon rank-sum test p-value < 4.8e-6; *controlled* vs *controlling* proteins). The *controlled* proteins also show a smaller size (Wilcoxon rank-sum test p-value < 9.8e-6; *controlled* vs *controlling* proteins), which corroborates the hypothesis that protein size is important for the buffering mechanism (**Figure 2E**). These results increased the evidence of interactions and regulators that may act as drivers of protein complex assembly.

**Figure 2.**
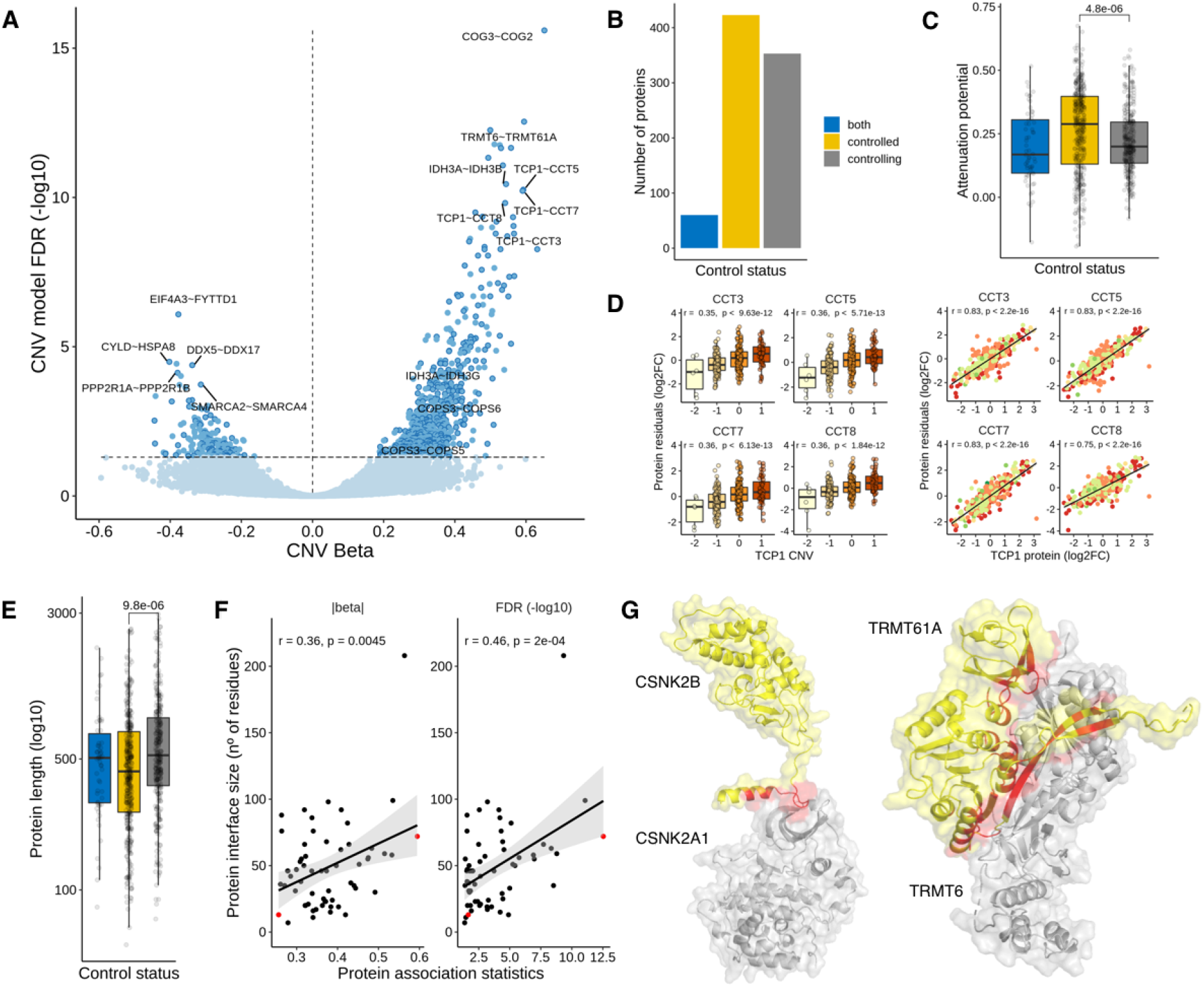
Physical protein associations. **(A)** Volcano plot of CNV beta (x-axis) and FDR (y-axis) for 411,591 protein pairs. Non-significant associations (FDR > 5%) are represented in light-blue, and significant associations (FDR < 5%) in dark blue. Associations also found to be significant (FDR < 5%) in the mRNA model and filtered by genomic co-localization are highlighted with a darker border (516). **(B)** Number of proteins by control status. **(C)** Distribution of attenuation potential by control status. **(D)** Examples of protein associations between TCP1 (*controlling* protein) and CCT3, CCT5, CCT7 and CCT8 (*controlled* proteins). The boxplots show the relation between the CNV changes of TCP1 and the protein residuals (log2FC) of the interacting partners. The scatter plots show the same relation with the protein abundance of TCP1. **(E)** Protein length (log10 of number of residues) by control status. **(F)** Scatter plots displaying the correlation between the protein association statistics (beta and FDR) with the protein interface size (number of residues at the protein interface, measured in the *controlled* protein). Each dot is a protein association. Two representative associations between CSNK2A1 - CSNK2B (small interface) and TRMT6 - TRMT61A (big interface) are denoted in red. **(G)** Representation of protein interactions between CSNK2A1 and CSNK2B and TRMT6 and TRMT61A. The *controlled* proteins are coloured in yellow (CSNK2B and TRMT61A) and the *controlling* proteins are coloured in *grey* (CSNK2A1 and TRMT6). The interface area is represented in red.

We hypothesized that protein interaction-dependent control of degradation could depend on the protein interfaces size. To test this, we identified 60 significant associations with available structural models (**Methods**) and correlated the protein interface size with the effect-size (beta value) and significance (FDR) of the respective protein association pairs (**Figure 2F**). We found that both statistics are positively and significantly correlated with interface size (CNV beta – Pearson’s r: 0.36; p-value: 4.5e-3; -log10 FDR – Pearson’s r: 0.46; p-value: 2.0e-4). We selected two examples to illustrate the observed differences (**Figure 2G**). Post-transcriptional regulation of TRMT61A by TRMT6, that form the tRNA (adenine-N1-)-methyltransferase enzyme, is the second strongest association found in our analysis, and the interface formed between these two proteins covers a total of 72 residues. In contrast, a weaker association between CSNK2A1 and CSNK2B may be explainable by a much smaller interface of 13 residues.

These results show that interface sizes are an important determinant of the protein interaction mediated control of protein degradation. This may be due to an effect of binding affinity or differences in the recognition of exposed interfaces of different sizes by the degradation machinery.

### Identification of phosphorylation sites that may modulate protein complex assembly

The role of phosphorylation in modulating protein binding affinities has been well described (Beltrao et al., 2012; Betts et al., 2017; Nishi, Hashimoto, & Panchenko, 2011). We reasoned we could use the multi-omics datasets to find protein interactions affected by phosphorylation, which in turn could impact complex assembly and protein degradation. Out of 368 samples with CNV, mRNA and protein measurements, 170 also have quantifications at the phosphosite level (**Figure 3A**). For this analysis we used proteins and phosphosites measured in at least 50% of the 170 samples, corresponding to 8,546 proteins and 5,733 phosphosites.

**Figure 3.**
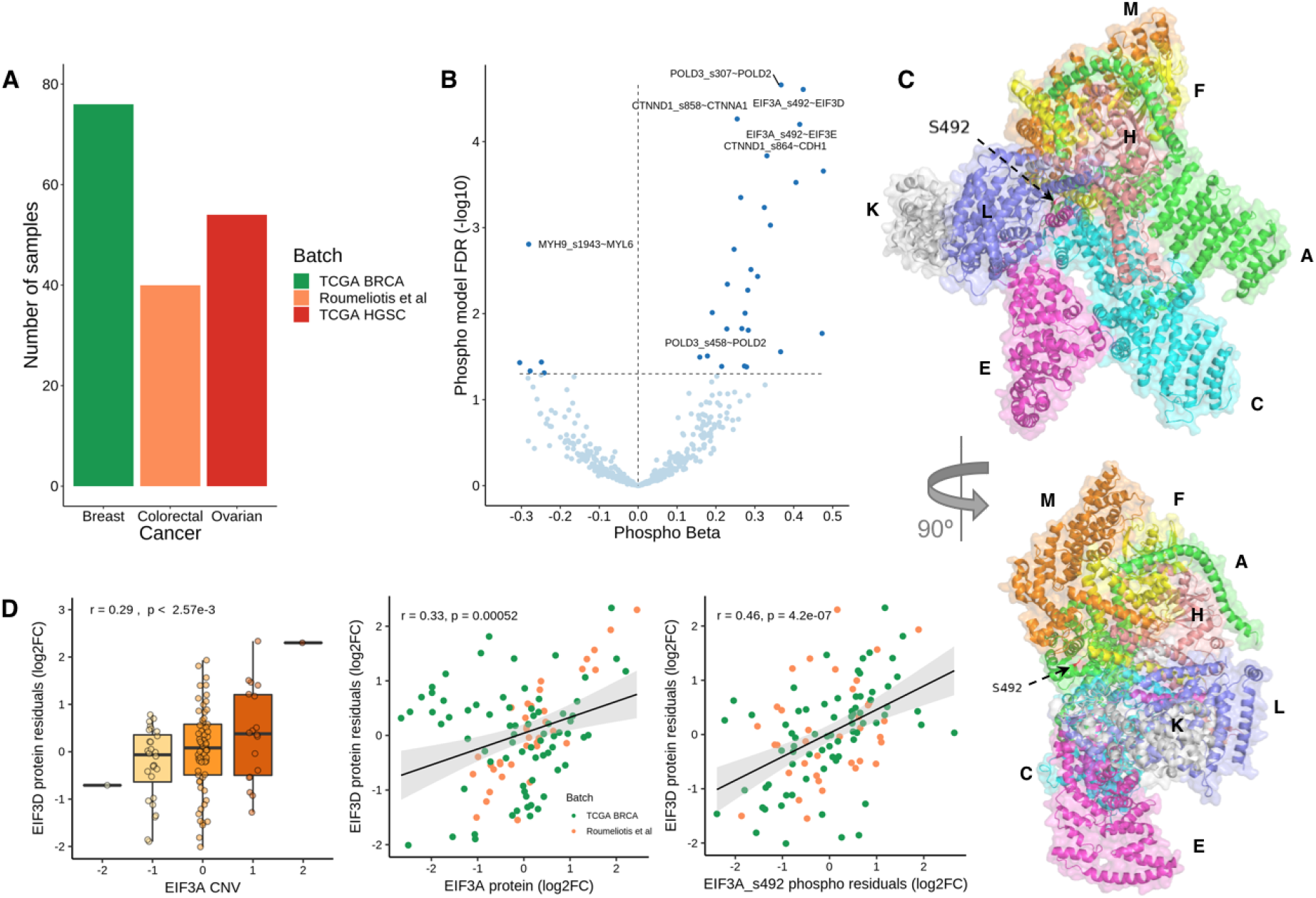
Identification of phosphorylation sites with a potential role in regulating protein interactions. **(A)** Number of samples with CNV, mRNA and phospho(protein) measurements, by cancer type/batch. **(B)** Volcano plot of phospho beta (x-axis) and FDR (y-axis). Each dot is a phosphosite-protein association, between a putative regulatory phosphosite Xp and a regulated protein Y. All associations (438) are significant in the CNV and mRNA models, between the putative regulatory protein X and the regulated protein Y. 32 associations (FDR < 5%) are also significant in the phospho model (dark blue). **(C)** Representation of EIF3 complex in two orientations. The arrow points to the phosphosite S492 (serine 492) at EIF3A subunit. **(D)** Significant association between EIF3A/EIF3A S492 and EIF3D. The boxplots show the agreement between the CNV changes of EIF3A and the protein residuals (log2FC) of EIF3D. The scatter plots show the same relation with the protein and phosphosite (S492) abundances of EIF3A.

Using the compendium of physical interactions (572,856 protein interactions), we tested whether the changes of a phosphosite Xp from protein X is associated with the protein levels of the interacting protein Y. As before we used a linear regression model where the protein abundance of protein Y is predicted using the phosphosite levels of protein X (Xp), while taking into account the protein and CNV levels of protein X, the RNA of protein Y, and other covariates (**Methods**). Out of 315,772 phosphosite-protein pairs tested with this model, 11,672 associations were significant (FDR < 5%). To ensure the associations are directional, we overlapped these associations with the 516 protein-protein associations found with the CNV and mRNA models, identifying 32 overlapping associations (**Figure 3B**, listed in **Supplementary Table 3**). Our interpretation of these associations is that these phosphosites can regulate the protein interaction and thereby modulate the degradation of the complex subunits.

The 32 associations involve 28 phosphosites, and of these 2 phosphosites are already known to regulate interactions (POLD3 S458 and MYH9 S1943) and an additional case (EIF3A S492) is not yet known to regulate protein interactions but is at the interface with other complex members (**Figure 3C**). EIF3A is predicted here to be a “rate-limiting” subunit of the eukaryotic initiation factor 3 complex and has been previously experimentally implicated in the control of protein levels of several of the other subunits (S. Wagner, Herrmannová, Malík, Peclinovská, & Valášek, 2014). One phosphosite of EIF3A (S492) showed a strong association with the protein levels of two other complex subunits (EIF3D and EIF3E). In line with this, we find that the copy number of EIF3A correlates with the residual protein levels of EIF3D (i.e. after regressing out EIF3D mRNA levels) and that the phosphosite levels of EIF3A S492 correlates better with EIF3D protein residual than the EIF3A total protein levels (**Figure 3D**). These results suggest that EIF3A S492 may have an impact on protein complex assembly.

### Protein attenuation mechanisms found in cancer are observed in normal tissues

The study of the impact of CNVs in cancer proteomes indicates that up to ~40% of genes have copy number changes that are buffered at the protein level. Such post-transcriptional regulatory processes should not be specific to cancer, however, the extent that these effects are observed in normal cellular states is still largely unknown. To address this question we analyzed gene and protein expression datasets for normal tissues, made available by the Genotype-Tissue Expression (GTEx) and Human Protein Map (HPM) projects. In total, we collected expression for 5,239 proteins and genes, across 14 tissue types (**Methods**).

We tested if the post-transcriptional control dependent on protein interactions observed in cancer is present in normal tissues. For this, we asked if the protein abundance of *controlling-controlled* protein pairs will tend to correlate more strongly than other protein interaction pairs. Similarly, we expected that the correlation between the mRNA and protein levels of *controlled* subunits would tend to be weaker than for non post-transcriptionally *controlled* proteins. We tested this using protein-protein interaction pairs measured in the tissue data with significant *controlling-controlled* relationships from cancer data (301 pairs) and all other 161,945 protein-protein interactions pairs (**Methods**). Reassuringly, we observed that the correlation of protein abundance across tissues increased for protein pairs with stronger association strength, for stable mRNA-mRNA correlation values (Wilcoxon rank-sum test p-value = 8.96e-4 between non-significant and significant pairs; p-value = 8.25e-06 between non-significant and highly-significant pairs) (**Figure 4A**). Also, as predicted the protein to mRNA correlation values across tissues, of the *controlled* subunits, decreases with the association strength (Wilcoxon rank-sum test p-value = 0.022 between non-significant and significant pairs) (**Figure 4A**). We provide two examples for protein interacting pairs ARCN1 and COPA, and TRAPPC8 and TRAPPC11 where the mRNA levels of the *controlling* subunits (ARCN1 and TRAPPC8) appear to dictate the protein abundance of both proteins (**Figure 4B**). These results suggest that the protein associations identified in the cancer datasets can also be observed in normal tissues, at least in aggregate. Importantly, they demonstrate that cancer data can be a useful resource to study protein homeostasis in normal conditions.

**Figure 4.**
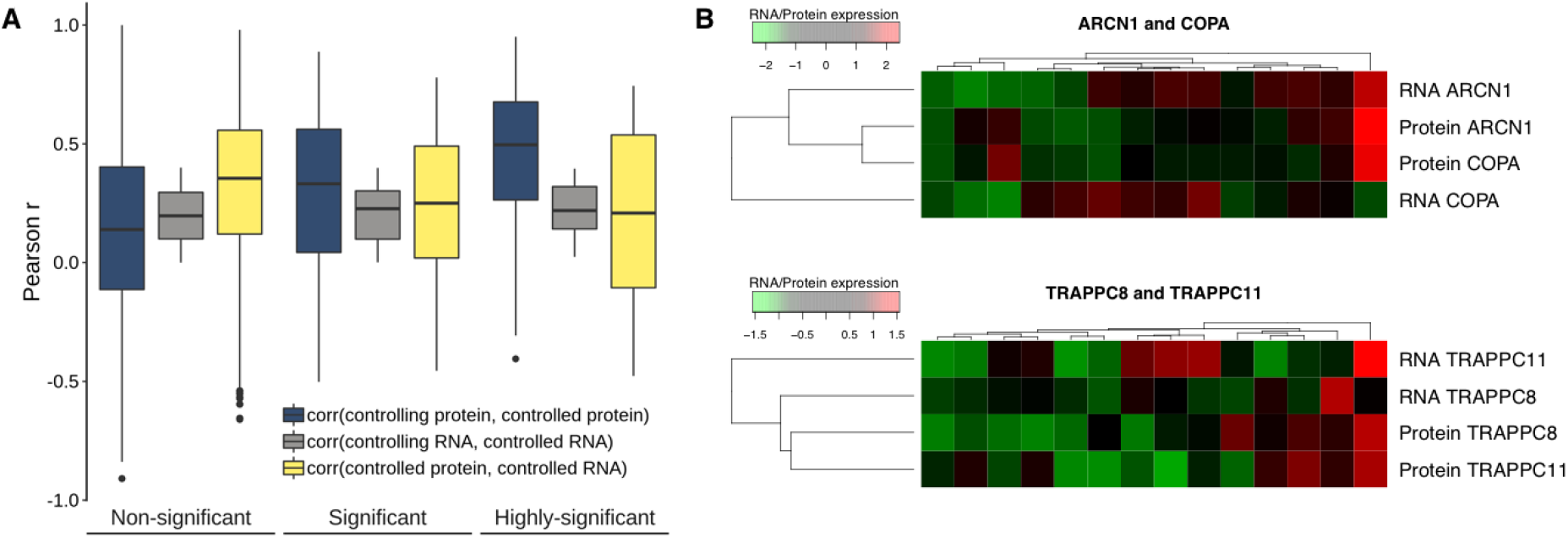
Evidence of interaction mediated control of protein abundances in normal tissues. **(A)** Pearson correlation coefficient between the protein of the *controlling* and *controlled* genes (blue); mRNA of the *controlling* and *controlled* genes (grey) and mRNA and protein abundance of the *controlled* gene (yellow); for the non-significant associations (FDR > 5%), significant associations (1% < FDR < 5%) and highly-significant associations (FDR < 1%). **(B)** Heatmap showing the agreement between the mRNA and protein expression profiles (rows) across tissues (columns) for two highly-significant associations: ARCN1 (*controlling*) ~ COPA (*controlled*) and TRAPPC8 (*controlling*) ~ TRAPPC11 (*controlled*).

### Buffering of gene expression variation due to natural genetic variation

If the phenomenon of interaction mediated control of protein abundances is important for tuning protein levels in normal cells, we then expect consequences on how natural variation may sometimes result in changes in mRNA but not protein and consequently phenotypic traits. In the context of the attenuation models studied here, single nucleotide polymorphisms (SNPs) associated with gene expression via quantitative trait loci (QTL) analysis – known as expression QTLs (eQTLs) – should also tend to be attenuated at protein level potentially for the same genes as those found in cancer. To study this, we focused on a reduced set of genes with significant CNV-mRNA correlation (Pearson’s r > 0.3), and analysed if protein level CNV buffering could affect the probability of eQTLs to have phenotypic impact and be associated with disease traits, as measured by tagging to SNPs linked to phenotypes in genome-wide association studies (GWAS) (**Figure 5A** and **Methods**). To this end we relied on cis-eQTLs reported in GTEx and compared the fraction of GWAS tagging eQTLs for different classes of protein attenuation (**Figure 5B top** and **Methods**). We found that eQTLs corresponding to genes classified as highly attenuated have a lower fraction of GWAS tagging eQTLs, and that the difference between the degree of attenuation increases for eQTLs mapped in multiple tissues (**Figure 5B top**). A logistic model linking the GWAS tagging status of the eQTLs to the attenuation score of the corresponding *cis* genes confirms that variation with impact on expression of attenuated proteins will tend to be buffered and have a lower chance of causing a phenotypic effect (**Figure 5B bottom**).

**Figure 5.**
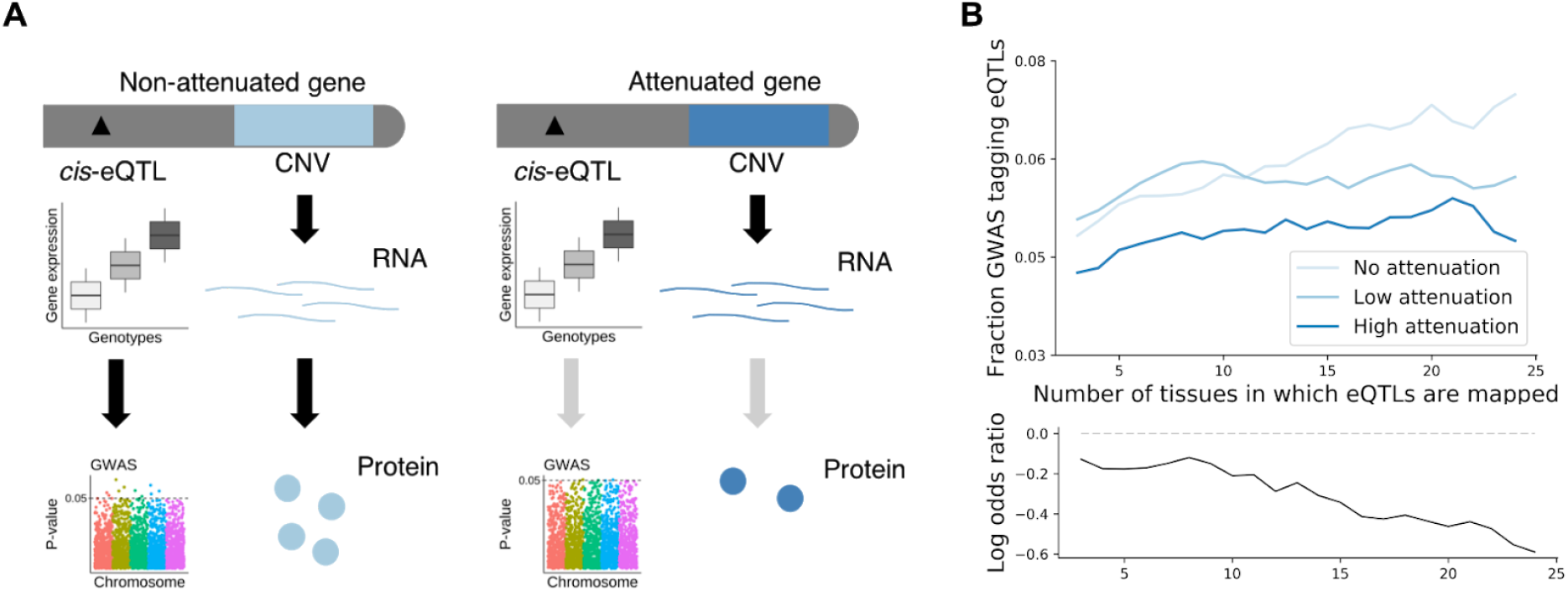
Protein attenuation reduces cis-eQTLs impact on phenotypic traits. **(A)** Diagram illustrating the highest probability of cis-eQTLs from non-attenuated genes (red) tag GWAS hits, in comparison to cis-eQTLs from attenuated genes (blue). **(B)** (top) Fraction of eQTLs associated to disease traits for the three classes of CNV attenuation: no (light-blue), low (blue) and high (dark-blue) attenuation. The x-axis corresponds to the number of tissues in which an eQTL was called. (bottom) Log odds ratio of eQTL association with a disease trait. Values indicate the change in the odds ratio of eQTL association with GWAS variants for an unit increase in the attenuation score.

Highly attenuated genes tend to be enriched in protein complexes that are more essential to the cell and therefore could have specific biases as to how eQTLs are linked to GWAS associated traits. To account for this potential bias we replicated the analysis on members of protein complexes. Interestingly, this shows that the attenuation score has a higher impact on GWAS tagging probability for members of protein complexes, and more specifically for members of large protein complexes (>5 subunits) (**Figure S5**).

These results suggest that the CNV attenuation measured in cancer cells for protein abundance has direct application in the ranking of potential impact of mRNA variation on phenotype differences and support the idea that some of these attenuation mechanisms may take place in multiple tissues.

## Discussion

The joint analysis of multi-omics datasets of cancer samples suggests that a very significant fraction of the proteome (up to 42%) is under post-transcriptional control. The set of genes with protein level buffering of CNVs is enriched in gene-products belonging to large protein complexes. In addition, we found that the fraction of interface residues of a protein is a strong determinant of attenuation. Together with experiments on pulse chase degradation (McShane et al., 2016), aneuploidy (Dephoure et al., 2014; Pavelka et al., 2010; Stingele et al., 2012) and the impact of natural genetic variation on protein levels (Battle et al., 2015; Chick et al., 2016) these results implicate protein complex formation as an important factor in post-transcriptional control, most likely via a high degradation rate of unassembled subunits. However, it is likely that multiple mechanisms contribute to post-transcriptional control measured in the cancer samples including, for example, the control of protein translation rates by microRNAs or RNA binding proteins. The extent of post-transcriptional control that is explained by the different processes remains to be studied.

We observed that the fraction of residues at the interface correlates with the probability that a protein shows gene dosage attenuation. Similarly, the size of the interface correlates with the strength of association between pairs of physical interactions in which one subunit appears to control the abundance level of the interaction partner. The size of the interface typically correlates with increasing binding affinity between proteins as well as larger amounts of hydrophobic residues that are exposed in the absence of interactions. We speculate that either of these consequences could play a role in the attenuation. In particular, larger fraction of hydrophobic regions could increase the propensity to form aggregates and in some cases hydrophobic regions are known to be recognized for degradation (Xu, Anderson, & Ye, 2016). This could represent a general mechanism for recognition of unassembled complex subunits. The structural analysis performed here is limited by the current lack of coverage for structures of protein complexes. In the future, additional structures may allow us to study in more detail the interface features that are important for the attenuation mechanism.

We have used data from cancer samples to identify the attenuated proteins and physical interactions with “rate-limiting” subunits. It is still unclear if the same proteins and interactions will have the same post-transcriptional control in other systems and/or species. When studying expression variation in normal tissues and the association of eQTLs with phenotypes we observed that, in aggregate, the same proteins and interactions show signals consistent with post-transcriptional buffering of mRNA expression variation. Of note, we find that eQTLs are less likely to be linked to phenotypes in highly attenuated proteins. This is in line with studies of mRNA and protein QTLs in human iPSC lines, showing that genetic variation driving mRNA changes are more likely to be associated to genotype differences when they are observed at the protein level (Mirauta et al., 2018). These findings highlight the importance of studying the degree of conservation of these post-transcriptional processes in different tissues and systems in the context of human genetics and disease.

## Methods

### Multi-omics data collection

Proteomics and phosphoproteomics quantifications at the protein/phosphosite level from TCGA cancer patients were obtained from the CPTAC data portal (proteomics.cancer.gov/data-portal), for breast cancer (BRCA) (Mertins et al., 2016), colorectal cancer (COREAD) (B. Zhang et al., 2014) and ovarian cancer (HGSC) (H. Zhang et al., 2016). The same data from cancer cell lines were downloaded for COREAD cell lines (Roumeliotis et al., 2017) and for BRCA cell lines (Lapek et al., 2017; Lawrence et al., 2015). Gene-level RNA-seq raw counts were acquired from GEO (GSE62944) (Rahman et al., 2015) for TCGA samples and from the CCLE data portal (Barretina et al., 2012; Cancer Cell Line Encyclopedia Consortium & Genomics of Drug Sensitivity in Cancer Consortium, 2015; Rahman et al., 2015) for cancer cell lines. Copy-number variation GISTIC levels (Mermel et al., 2011) were compiled from the firebrowse (firebrowse.org) data portal (accession date 15/01/2018) for TCGA samples and from the CCLE data portal for cancer cell lines (accession date 14/02/2017).

### Data pre-processing and normalisation

The label-free protein quantifications (precursor areas) for COREAD CPTAC samples (B. Zhang et al., 2014) were first normalized by sample, where summed peak areas for the same protein were divided by the total summed area for the observed sample proteome. Relative protein abundances were then calculated by dividing each protein area over the median area across samples, and then log2 transformed. Protein and phosphosite intensities for COREAD cell lines (Roumeliotis et al., 2017) were divided by 100 and transformed to log2. For BRCA cell lines (Lapek et al., 2017) protein log2 fold-changes were calculated by subtracting the median intensities across the samples. Similarly, the label-free protein intensities (peak areas) for BRCA cell lines from (Lawrence et al., 2015) were converted into relative abundances by calculating the log2 ratio of protein intensities over the median intensities across samples. Sample replicates of protein and phosphoprotein were combined by averaging the values for each protein and phosphosite, respectively. Phosphopeptides intensities mapping to the same phosphosite were combined by calculating the median phosphosite intensity per sample. In the cancer cell lines, genes with multiple isoforms were filtered by selecting the protein isoform with highest median expression across samples. Proteomics and phosphoproteomics distributions across cancer samples and cell lines were quantile normalized to ensure comparable distributions, using *normalizeQuantiles* function from Limma R package (Ritchie et al., 2015). In total, 13,569 proteins across 436 samples (340 cancer samples and 96 cell lines) and 79,824 phosphosites across 195 samples (145 cancer samples and 50 cell lines) were assembled in this study. Given the sparseness of the phospho(protein) data, for the subsequent analyses it was only selected proteins measured in at least 25% of the 368 samples with protein, mRNA and CNV measurements, and the phosphosites measured in at least 50% of the 170 samples with also phosphorylation data, comprising 9,188 proteins and 5,733 phosphosites. The phospho(protein) and mRNA data were then standardized using the z-score transformation.

At the RNA-seq level, lowly expressed genes were removed by filtering out genes with mean counts-per-million (CPM) lower than 1 across samples. After raw counts normalization by the trimmed-mean of M-values method (Robinson & Oshlack, 2010) using the edgeR R package (Robinson, McCarthy, & Smyth, 2010), the log2-CPM values were extracted from *voom* (Law, Chen, Shi, & Smyth, 2014) function in Limma. After merging the CPTAC samples with the CCLE cell lines, the final RNA-seq dataset comprised 13,228 genes with measurements across 370 samples (296 cancer samples and 74 cell lines). At the CNV level, after compiling the GISTIC thresholded data, 19,023 genes were found to have CNV measurements across 412 samples (337 cancer samples and 75 cell lines).

Potential confounding factors revealed by PCA analysis (supplementary figures 1A and 2A) were regressed-out using a multiple linear regression model. This model was implemented with the protein or mRNA abundance of a given gene as dependent variable and the potential confounding factors, i.e. cancer type, experimental batch, proteomics technology, age and gender as independent variables. The residuals from the linear model were the protein and mRNA variation not driven by the confounding effects, as the second PCA demonstrated (supplementary figures 1B and 2B).

### Analysis of protein attenuation

The strategy in (Gonçalves et al., 2017) was used to evaluate the impact of copy-number variations at the genome level on cancer proteomes. For each gene, the Pearson correlation coefficients between the CNV and mRNA and the CNV and protein were calculated, and an attenuation measure devised as follows:

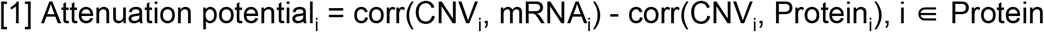

where *corr* represents the Pearson correlation coefficient and *Protein* represents 8,124 genes for which CNV, mRNA and protein quantifications across 368 samples where available. After calculating the attenuation potentials, a gaussian mixture model (GMM) with 4 mixture components was used to cluster the genes in four different groups. Group 1 had 19 genes with a negative attenuation potential, due to the higher correlation between the CNV and Protein than with the CNV and mRNA. These genes, which were not attenuated at the protein level, were included with the remaining non-attenuated genes in group 2, comprising 4,689 genes. Groups 3 and 4 contained the lowly-attenuated and highly-attenuated genes, with 2,578 and 857 genes, respectively. The GMM was implemented using *Mclust* function from the mclust R package (Scrucca, Fop, Murphy, & Raftery, 2016).

The enrichment of CORUM complexes was calculated with an hypergeometric test, using the *enrichr* function from the *clusterProfiler* R package (Yu, Wang, Han, & He, 2012). Only CORUM complexes with a Jaccard index lower than 0.9 and with more than 5 proteins were used. The comparison of ubiquitination sites fold-changes across protein attenuation levels was done using protein ubiquitination data obtained with three proteasome inhibitors: MG-132, epoxomicin and bortezomib (Higgins et al., 2015; W. Kim et al., 2011; Udeshi et al., 2013; S. A. Wagner et al., 2011).

### Compendium of physical protein interactions

In order to build a compendium of physical protein interactions, we downloaded a data set of protein-protein interactions from BioGRID version 3.4.157 (Stark et al., 2006) (accession date 30/01/2018). We only selected protein interactions occurring in human and captured with physical experimental systems. Interactions captured with Affinity Capture-RNA and Protein-RNA were excluded in order to guarantee that our dataset contained only interactions observed at the protein level. After excluding protein homodimers, 524,148 protein interactions (262,074 unique) were compiled with BioGRID. A list of protein interactions was also built using a set of protein complexes from the CORUM database (Giurgiu et al., 2018) (accession date 29/05/2018). The rationale was that protein partners from the same protein complex interact physically at least once. Using a set of 1,787 protein complexes and excluding protein homodimers, we assembled 74,712 (37,356 unique) physical protein interactions. This was expanded using a set of curated protein complexes from the endoplasmic reticulum (ER), yielding 1,196 (598 unique) protein pairs. In total, 572,856 (286,428 unique) protein physical interactions were compiled.

### Linear modelling to identify protein and phospho-protein associations

#### Protein associations

For a given protein physical interaction pair X and Y, it was tested whether protein X can control the protein levels of Y through protein-protein interaction, potentially constraining the degradation rate of Y. For each interacting pair two nested linear models were fitted. The first model (null) was used to predict the protein levels of Y (Py) using its mRNA (Ty) and a set of other covariates, i.e. cancer type, experimental batch, proteomics technology, patient age and gender (equation 1). In a second linear model (alternative), the CNV levels of X (Gx) was added as predictor variable (equation 2). A Likelihood Ratio Test (LRT) (equation 3) was then applied, in order to test whether the second model increases the goodness of fit of the first model in predicting Py.

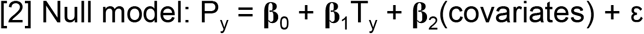

**β**_0_ represents the intercept, **β**_1_ the regression coefficient (effect size) for the mRNA of Y, **β**_2_ the regression coefficients of the covariates, and ε the noise term.

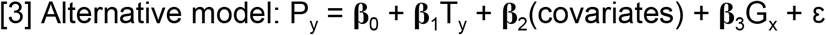

**β**_3_ is the regression coefficient for the CNV (G_x_) of protein X. A likelihood ratio test (LRT) was used to assess the significance of the association:

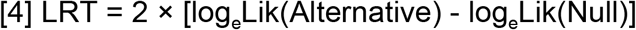

log_e_Lik corresponds to the log likelihood of the alternative and null models. P-values were then calculated using the LRT statistic over a chi-squared distribution, and adjusted for false discovery rate (FDR) using the Benjamin-Hochberg method. This model was applied for a given protein association pair X and Y if: X ∈ CNV Λ Y ∈ Protein Λ Y ∈ mRNA, where CNV, Protein and mRNA represent the multi-omics datasets.

A total of 411,591 protein pairs followed this criteria and were tested across 368 tumor samples. The same analysis was performed with the mRNA, instead of the CNV, of protein X for 392,128 protein pairs. To avoid spurious protein associations that might occur due to the genomic co-localization of the *controlling* proteins, the top ranked association was selected using the Borda ranking method. This was done systematically for every cases where multiple *controlling* proteins in the same chromosome were associated with the same *controlled* protein. More than one *controlling* protein in the same chromosome for the same *controlled* protein were allowed if their CNV profile Pearson correlation was lower than 0.5.

The linear models were implemented using *lm* R function and the LRT test with associated statistics were calculated using *Irtest* function from lmtest R package. The Borda ranking method was implemented using the *Borda* function from TopKLists R package (Schimek et al., 2015).

#### Phospho-protein associations

For a given protein pair X and Y it was tested whether a phosphosite Xp from protein X can be associated with changes in the protein abundance of protein Y. A similar model to before linear regression models and LRT tests was used. For each phosphosite-protein interaction, a first null model was fitted to predict the protein levels of Y (Py) using its mRNA (Ty), the CNV and protein levels of protein X (Gx and Px) and the same covariates (cancer type, patient age and gender) (Equation 4). In a second alternative linear model, the phosphosite Xp (Phox) of protein X was added as predictor variable (equation 5). The models were then compared using a Likelihood Ratio Test as in equation 3.

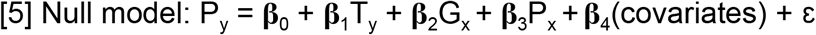

where **β**_0_ represents the intercept, **β**_1_ the coefficient of the mRNA of Y, **β**_2_ and **β**_3_ the regression coefficients for the CNV and Protein of X, respectively, and **β**_4_ the coefficients of other covariates, and ε the noise term.

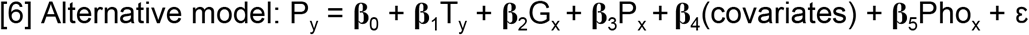

where **β**_5_ is the regression coefficient for the phosphosite Xp of protein X. This model was applied for a given phosphosite-protein association pair Xp and Y if Xp ∈ Phospho Λ X ∈ Protein Λ X ∈ CNV Λ Y ∈ Protein Λ Y ∈ mRNA, where Phospho, Protein, CNV, and mRNA represent the multi-omics datasets. A total of 315,772 phosphosite-protein pairs followed this criteria and were tested with this model across 170 tumor samples.

### Structural analysis

Protein interface sizes were calculated using an in-house pipeline (int3dInterfaces, github.com/evocellnet/int3dInterfaces) that extracts protein interfaces from Interactome3D structures (Mosca et al., 2012). For each protein interaction structure in Interactome3D, this pipeline uses NACCESS (bioinf.manchester.ac.uk/naccess) to calculate the solvent accessibility of the bound and unbound monomers. Every residue changing its relative solvent accessibility is considered to form part of the interface. From the 11,530 human protein interaction structures analysed with this pipeline, structures of protein homodimers or structures with less than 100 amino-acids were removed. Also, structures with chain lengths bigger than the respective UniProt protein lengths and with the same chain length for each partner were removed. After applying these filters, 3,082 structures with 6,147 protein interactions were used in the subsequent analyses.

For the 1,470 proteins which contained both information about CNV attenuation and interface size, the percentage of residues in protein interfaces was calculated as the ratio of the number of unique residues in interfaces over the protein size. For 60 significant protein association pairs represented in the structural data, the relation between the protein interface size with the regression CNV coefficient and FDR, was assessed using the Pearson correlation coefficient. For each pair, the protein interface size was calculated in the *controlling* and *controlled* proteins. The protein sizes (number of residues) were obtained from UniProt for 20,349 proteins (accession date 19/06/2018).

The percentage of area inside complex for the protein subunits from the COP9 signalosome was calculated using FreeSASA (Mitternacht, 2016). For each protein subunit, this percentage corresponded to the difference between the solvent accessible surface area (SASA) outside and inside complex over the SASA outside complex. The SASA was calculated in units of squared Ångström (Å^2^).

### Analysis of gene essentiality using CRISPR-Cas9 screenings

Gene essentiality data obtained with CRISPR-Cas9 screenings (Meyers et al., 2017) was downloaded from Project Achilles data portal (portals.broadinstitute.org/achilles) (accession date 31/10/2017). This data contains gene-dependency levels adjusted for copy-number specific effects for 17,670 genes across 341 cancer cell lines. Genes with an essentially score lower than -1×SD (the standard deviation for the entire data set corresponds to 0.3) in more than 5% of the cell lines were considered essential, and used in the remaining analysis (5,532 genes). The median gene essentiality was calculated for 3,548 genes with attenuation and essentiality data across the 341 cancer cell lines.

### Pairwise correlation of protein association pairs using normal tissue data

Gene and protein expression data for normal human tissues were obtained from the GTEx (GTEx Consortium et al., 2017) and Human Proteome Map (HPM) (M.-S. Kim et al., 2014) portals. The gene expression was obtained in the format of RNA-seq median RPKM for 56,238 genes across 53 tissues. The protein expression was downloaded as averaged label-free spectral counts for 17,294 genes across 30 tissues. For the protein expression data, it was selected 9,156 genes in common with the HPM data available in Expression Atlas (Petryszak et al., 2014). The 14 tissues common to the GTEx and the HPM used in the remaining analysis were: frontal cortex, spinal cord, liver, ovary, testis, lung, adrenal gland, pancreas, kidney, urinary bladder, prostate gland, heart, esophagus and colon. The gene expression in the last three tissues was averaged in GTEx, between heart atrial appendage and left ventricle; between esophagus gastroesophageal junction, mucosa and muscularis; and between colon sigmoid and transverse. The protein and gene expression data was then filtered to only include genes and proteins expressed in at least 10 of 14 tissues, resulting in 5,239 genes consistently expressed at the gene and protein level. The RNA and protein measurements were then standardized to z-scores and quantile normalized.

Having assembled the gene and protein expression datasets for normal tissues, pairwise Pearson correlation coefficients were calculated between the protein of the *controlling* and *controlled* genes, mRNA of the *controlling* and *controlled* genes, and mRNA and protein of the *controlled* gene. The Pearson correlations were calculated for 91 highly-significant associations (FDR < 0.01), 210 significant associations (0.01 ≤ FDR < 0.05) and 161,945 non-significant associations at the CNV and mRNA level (FDR ≥ 0.05).

### Analysis of the impact of CNV attenuation on the eQTL association to disease traits

Following the approach in HipSci proteomics (Mirauta et al., 2018), we considered a stringent set of 21,601 associations from the NHGRI-EBI GWAS catalog (download on 10 April 2018; converted to hg19) for analysis. We considered eQTLs reported from the Genotype-Tissue Expression (GTEx) in 35 tissues (excluding brain), compute the number of tissues having the same slope sign, i.e direction of effect size, and discarded those with consistent slope in less than 3 tissues.

We defined proxy variants of each cis-eQTL as variants in high LD (r^2^ > 0.8; based on the UK10K European reference panel) within the same *cis* window. Next we grouped eQTLs in high LD blocks (r^2^ > 0.8), excluded from this analysis 247 genes having each more than 100 eQTL blocks, and obtain a final set of 66,197 eQTL blocks corresponding to 2,953 genes and 441,194 eQTL – gene associations. We then define these blocks as GWAS-tagging if for at least one eQTL in the block at least one LD proxy variant was annotated in the NHGRI-EBI GWAS catalog. Finally, we report the fraction of GWAS-tagging eQTL stratified by the attenuation level of the corresponding *cis* genes. To assess the robustness of this analysis and to study the effects on GWAS tagging probability of eQTL recurrence across tissues, we compute the number of tissues in which an eQTL was called with the same slope, and report results by stratifying the eQTLs by increasing number of tissues.

We rely on core protein complexes from CORUM to identify the gene complex membership status, and segregate those which are annotated in at least one large complex (>5 subunits). Out of the genes with eQTL evidence and with annotation scores, 961 are annotated in CORUM and 576 are members of large complexes.

## Supplementary Figures and Tables

**Supplementary figure 1.**
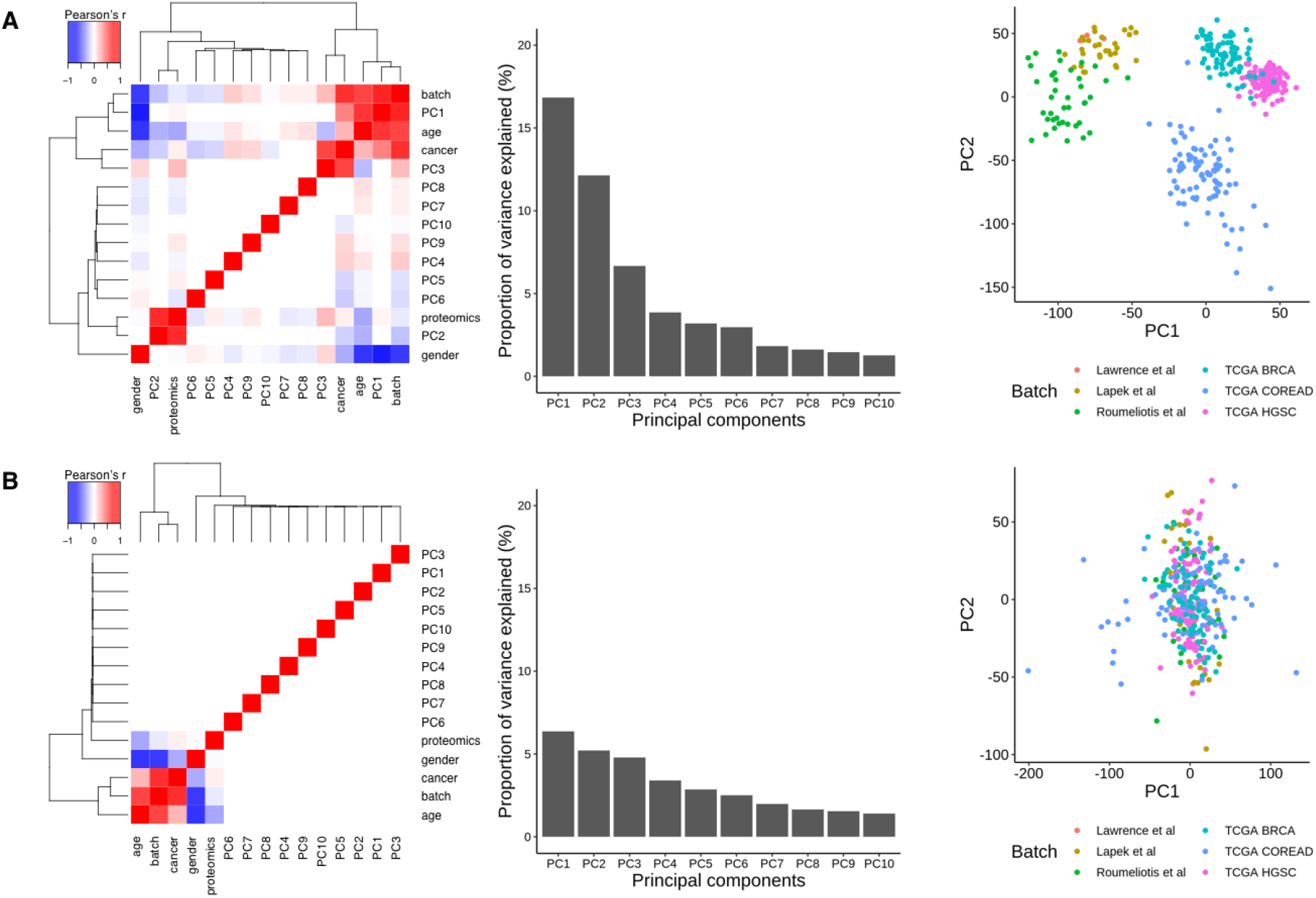
Confounding effects regressed-out from transcriptomics data. **(A)** Pearson correlation coefficient of the first 10 principal components (PCs) with the potential confounding effects before normalization. **(B)** Pearson correlation coefficient of the first 10 principal components (PCs) with the potential confounding effects after normalization.

**Supplementary figure 2.**
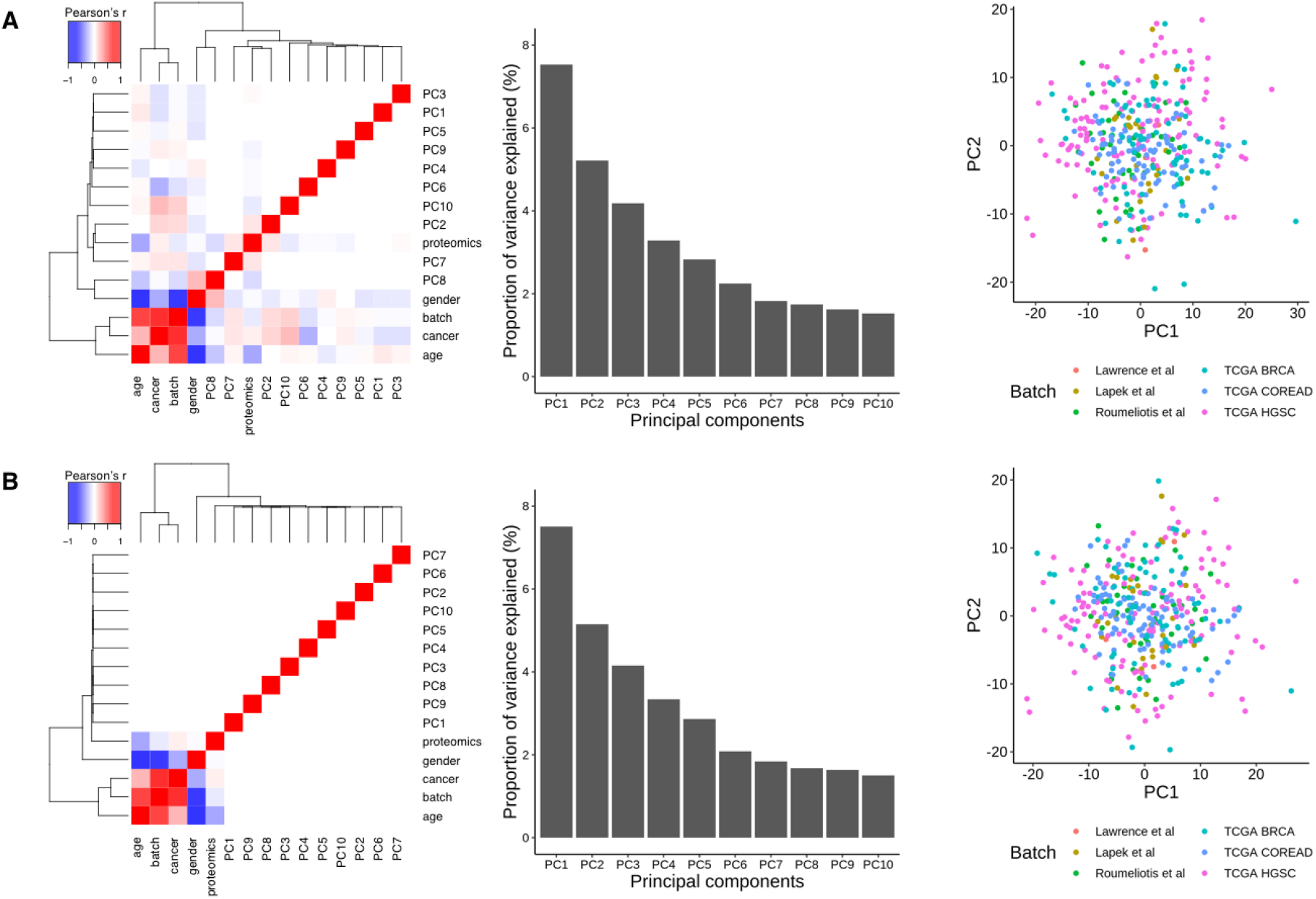
Confounding effects regressed-out from proteomics data. **(A)** Pearson correlation coefficient of the first 10 principal components (PCs) with the potential confounding effects before normalization. **(B)** Pearson correlation coefficient of the first 10 principal components (PCs) with the potential confounding effects after normalization.

**Supplementary figure 3.**
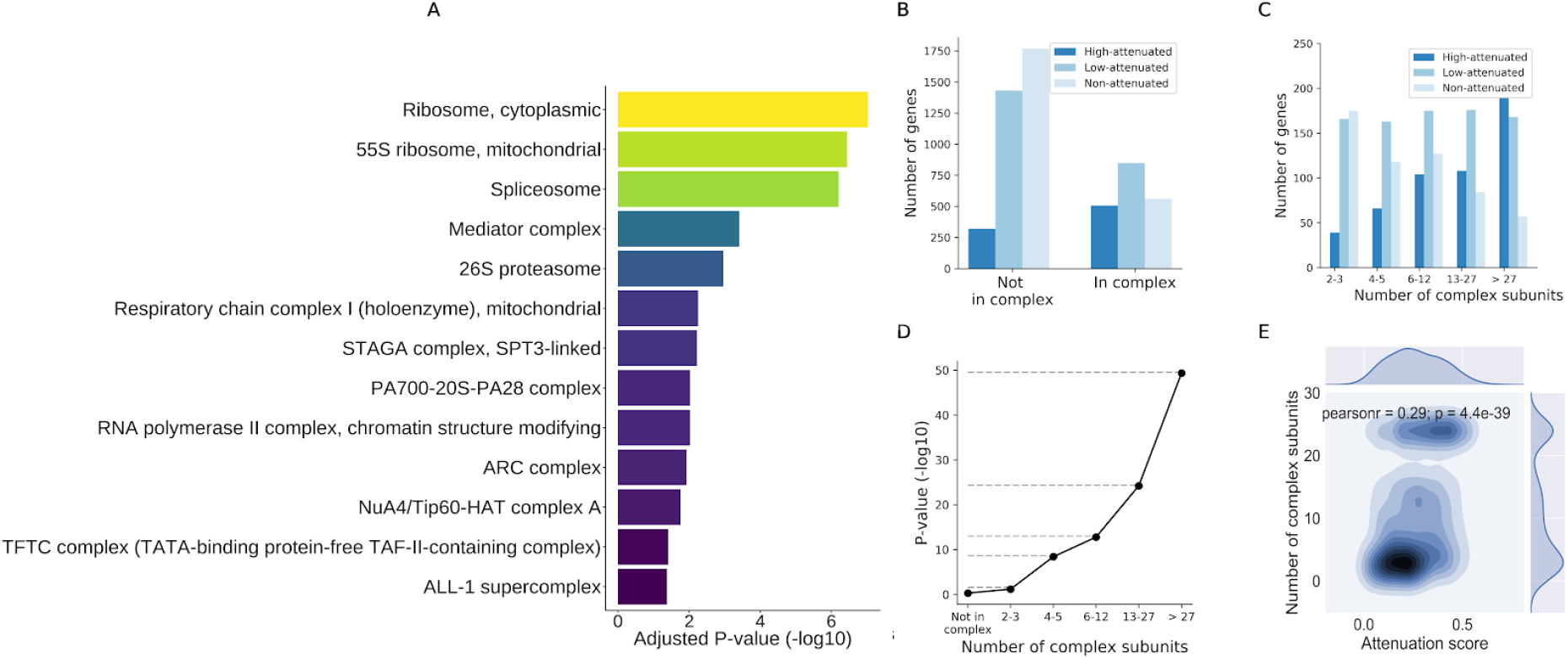
Relationship between CNV attenuation in proteins and protein complex membership. **(A)** List of top protein complexes ordered by enrichment in attenuated genes. X-axis shows p-values derived with an hypergeometric test (Benjamini-Hochberg multiple test correction). We discarded from the list of CORUM complexes those with a Jaccard index higher than 0.9 with any other complex and those with 5 proteins or less **(B)** Number of genes for each attenuation class by complex membership status (CORUM). **(C)** Number of genes stratified by the maximum number of subunits of any protein complex incorporating the genes. **(D)** Enrichment of attenuated proteins in members of protein complexes stratified by complex size, i.e number of subunits. Shown are the p-values derived with an exact Fisher test (alternative “greater”). **(E)** Relationship between the number of subunits in a protein complex and the protein complex member CNV attenuation. For each member of a complex we juxtapose the attenuation score (x-axis) and the maximum number of subunits of any complex this protein is part of (y-axis).

**Supplementary figure 4.**
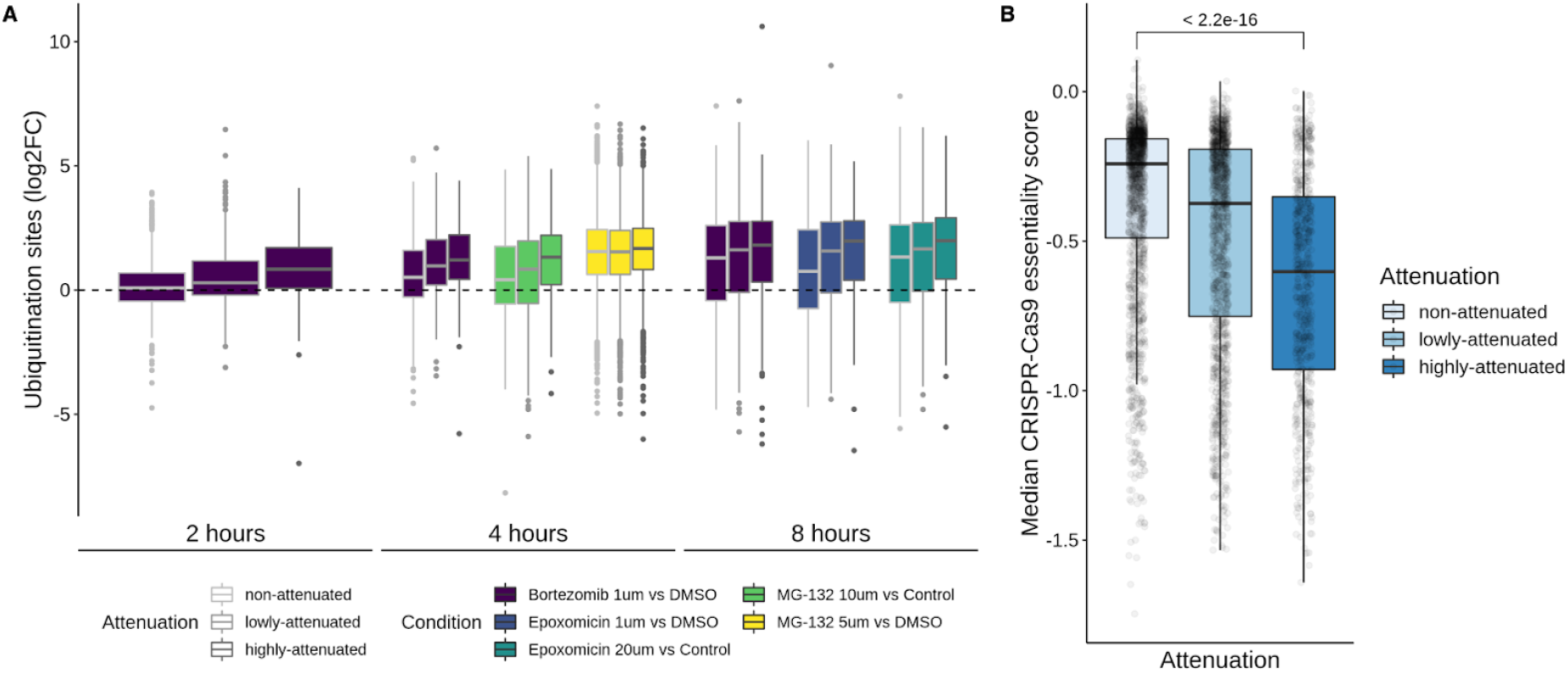
Attenuated proteins show faster increase in protein ubiquitination after proteasome inhibition and higher gene essentiality. **(A)** Ubiquitination sites fold-changes (y-axis) across protein attenuation levels (x-axis) after proteasome inhibition with three inhibitors: Bortezomib, Epoxomicin and MG-132. **(B)** Median gene essentiality measured in CRISPR-Cas9 screenings (y-axis), across 341 cancer cell lines, by attenuation level (x-axis).

**Supplementary figure 5.**
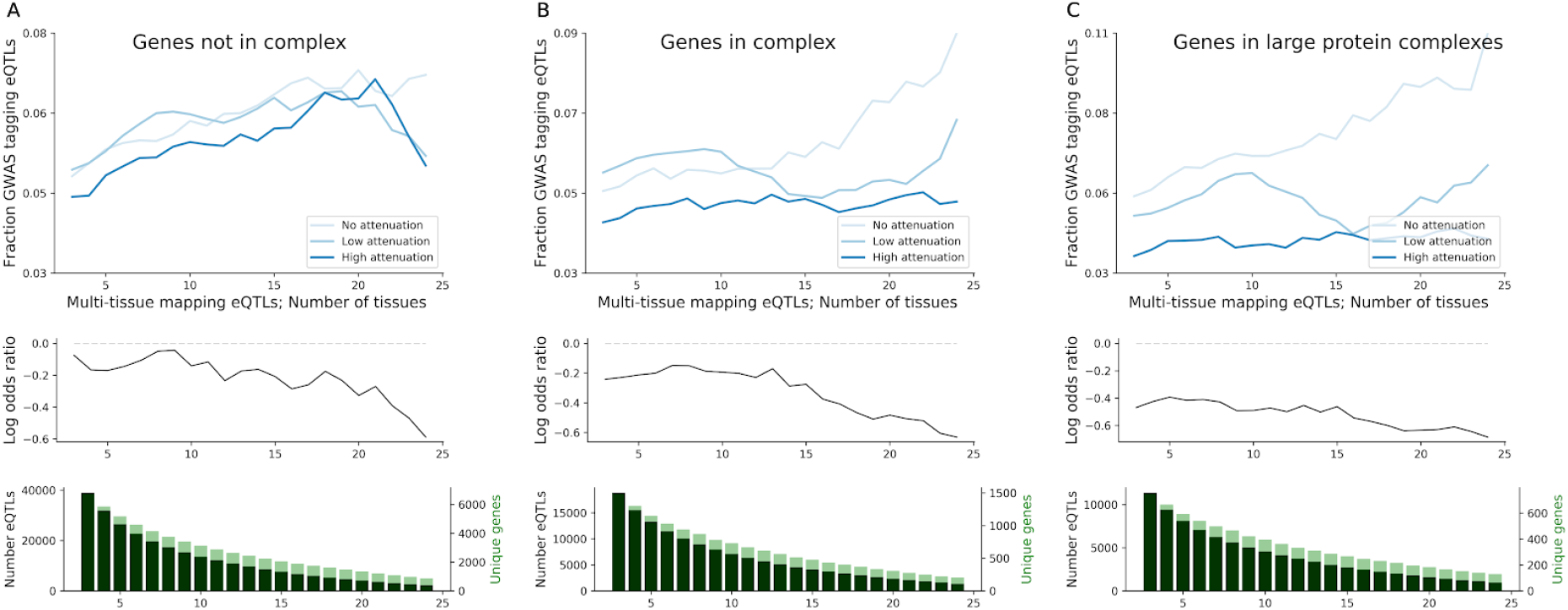
Analysis of the protein complex membership status of genes impact on the relationship between the CNV attenuation at protein level and eQTL association with disease traits. Identical analysis as in **Figure 3D** is performed for genes with no (A) and with (B) existing annotation in CORUM protein complexes, and for genes members of protein complexes with at least 5 subunits (C).

**Supplementary table 1.** 8124 genes stratified by attenuation level. The table includes the Pearson correlation coefficient between the CNV and mRNA and the CNV and protein, respective p-values and attenuation potential.

**Supplementary table 2.** 516 protein-protein associations significant in the CNV and mRNA models (FDR < 5%). For each association, the table includes the controlling and controlled protein, and the effect size (beta) and FDR from both models.

**Supplementary table 3.** 32 significant phospho-protein associations (FDR < 5%). The table includes the controlling protein/phosphosite, the controlled protein, and the effect size (beta) and FDR from the the phospho model. All associations are also significant in the CNV and RNA models, between the putative regulatory and regulated proteins.

## Supporting information

